# Custom built scanner and simple image processing pipeline enables low-cost, high-throughput phenotyping of maize ears

**DOI:** 10.1101/780650

**Authors:** Cedar Warman, John E Fowler

## Abstract

High-throughput phenotyping systems are becoming increasingly powerful, dramatically changing our ability to document, measure, and detect phenomena. Unfortunately, taking advantage of these trends can be difficult for scientists with few resources, particularly when studying nonstandard biological systems. Here, we describe a powerful, cost-effective combination of a custom-built imaging platform and open-source image processing pipeline. Our maize ear scanner was built with off-the-shelf parts for <$80. When combined with a cellphone or digital camera, videos of rotating maize ears were captured and digitally flattened into projections covering the entire surface of the ear. Segregating GFP and anthocyanin seed markers were clearly distinguishable in ear projections, allowing manual annotation using ImageJ. Using this method, statistically powerful transmission data can be collected for hundreds of maize ears, accelerating the phenotyping process.

## Introduction

High-throughput plant phenotyping is rapidly transforming crop improvement, disease management, and basic research (reviewed in Fahlgren et al., 2015; Mahlein, 2016; Tardieu et al., 2017). However, commercial phenotyping platforms remain out of reach for many laboratories, often requiring large initial investments of thousands or tens of thousands of dollars. While the cost of high-throughput sequencing has rapidly decreased, the cost of high-throughput phenotyping has remained high. New methods of low-cost, large-scale phenotyping are required to fully leverage the increasing availability of large datasets (e.g., genome sequences) and relevant quantitative statistical tools.

High-throughput phenotyping methods have been developed in several agricultural and model plant systems, including Arabidopsis and maize. Arabidopsis is well-suited to high-throughput phenotyping due to its small stature, rapid growth, and simple culture. Various systems have been created to measure Arabidopsis roots (Yazdanbakhsh and Fisahn, 2009; Slovak et al., 2014), rosettes (Arvidsson et al., 2011; Zhang et al., 2012; Awlia et al., 2016) and whole plants (Jiang et al., 2014). Most of these systems require robotic automation, which can drive up costs. Attempts at reducing costs rely on simple cameras and open-source image processing computational pipelines (Vasseur et al., 2018). Unlike Arabidopsis, maize plants are large, have a long growth cycle, and are typically grown seasonally outdoors. Because of these characteristics, maize is inherently more difficult to phenotype than Arabidopsis. However, as a consequence of its agricultural importance and utility as a model system, there has been substantial progress towards deploying maize phenotyping systems, both in the private (Choudhury et al., 2016) and academic (Miller et al., 2017) realms. As with Arabidopsis, several systems focus on phenotyping maize roots (Clark et al., 2013; Jiang et al., 2019) and above-ground vegetation (Chaivivatrakul et al., 2014; Junker et al., 2014; Choudhury et al., 2016; Zhang et al., 2017). Like with Arabidopsis, these systems are largely dependent on costly robotics and cameras.

Among maize tissues, ears are another target of interest for high-throughput phenotyping. Ears, with the seeds they carry, contain information about the plant and its progeny. They are easily stored, and do not require phenotyping equipment to be in place in the field or greenhouse at specific times during the growing season. Ears are a primary agricultural product of maize, which has led the majority of previous phenotyping efforts to focus on aspects of the ear that influence yield, such as ear size, row number, and seed dimensions (Liang et al., 2016; Miller et al., 2017; Makanza et al., 2018). These studies have used techniques that varied from expensive and specialized three-dimensional or line-scanning cameras (Wen et al., 2019; Liang et al., 2016) to relatively low-cost flatbed scanners and digital cameras (Miller et al., 2017; Makanza et al., 2018).

Beyond their agricultural importance, studying maize ears can answer fundamental questions about basic biology. The transmission of mutant genes can be easily tracked in maize seeds by taking advantage of a wide variety of visible endosperm markers (Neuffer et al., 1997; Li et al., 2013), which can be genetically linked to a mutant of interest (e.g. Arthur et al., 2003; Phillips and Evans, 2011; Bai et al., 2016; Huang et al., 2017). On the ear, seeds occur as an ordered array of progeny, which allows the transmission of mutant alleles to be tracked not only by individual cross, but within individual ears. The transmission of marker genes has thus far been quantified by hand, either by counting seeds on ears or after they have been removed. This approach has several limitations, among them a lack of a permanent record of the surface arrangement of seeds on the ear. The same disadvantages apply to most high-throughput seed phenotyping methods, which generally rely on seeds being removed from the ear before scanning and do not typically include marker information.

Here we address this missing aspect of high-throughput phenotyping in maize. Our rotational ear scanner and image processing pipeline is a cost-effective method for high-throughput ear phenotyping. By taking advantage of the cylindrical form of the maize ear, flat projections can be produced that provide a digital record of the surface of the ear, which can then be quantified in a variety of ways to track seed markers. Limiting materials to easily acquired parts and a basic camera makes this approach accessible to most if not all labs.

## Results

### Design and construction of the maize ear scanner

To efficiently phenotype maize ears, we designed a simple, custom-built scanner (Ear Rotational Scanner, ERS, v1.0) centered around a rotating ear. To scan the entire surface of the roughly cylindrical ear, the ear is rotated 360° while a stationary camera records a video, which can then be processed into a cylindrical projection. Materials for constructing the scanner were limited to those that were widely available and cost-effective (Table 1). The frame of the scanner was built from dimensional lumber, with a movable mechanism built from drawer slides that enables a wide range of ear sizes to be accommodated (Figure 1A). A rotisserie motor spins the ear at a constant speed, which is then imaged with a standard digital camera or cell phone (Figure 1B). The scanning process takes approximately 1 minute per ear, including the insertion of the ear into the scanner and video capture. For scanning ears carrying an engineered GFP marker that is highly expressed in the endosperm (Li et al., 2013), ears were illuminated with a blue LED light with an orange filter placed in front of the camera.

**Table 1.**
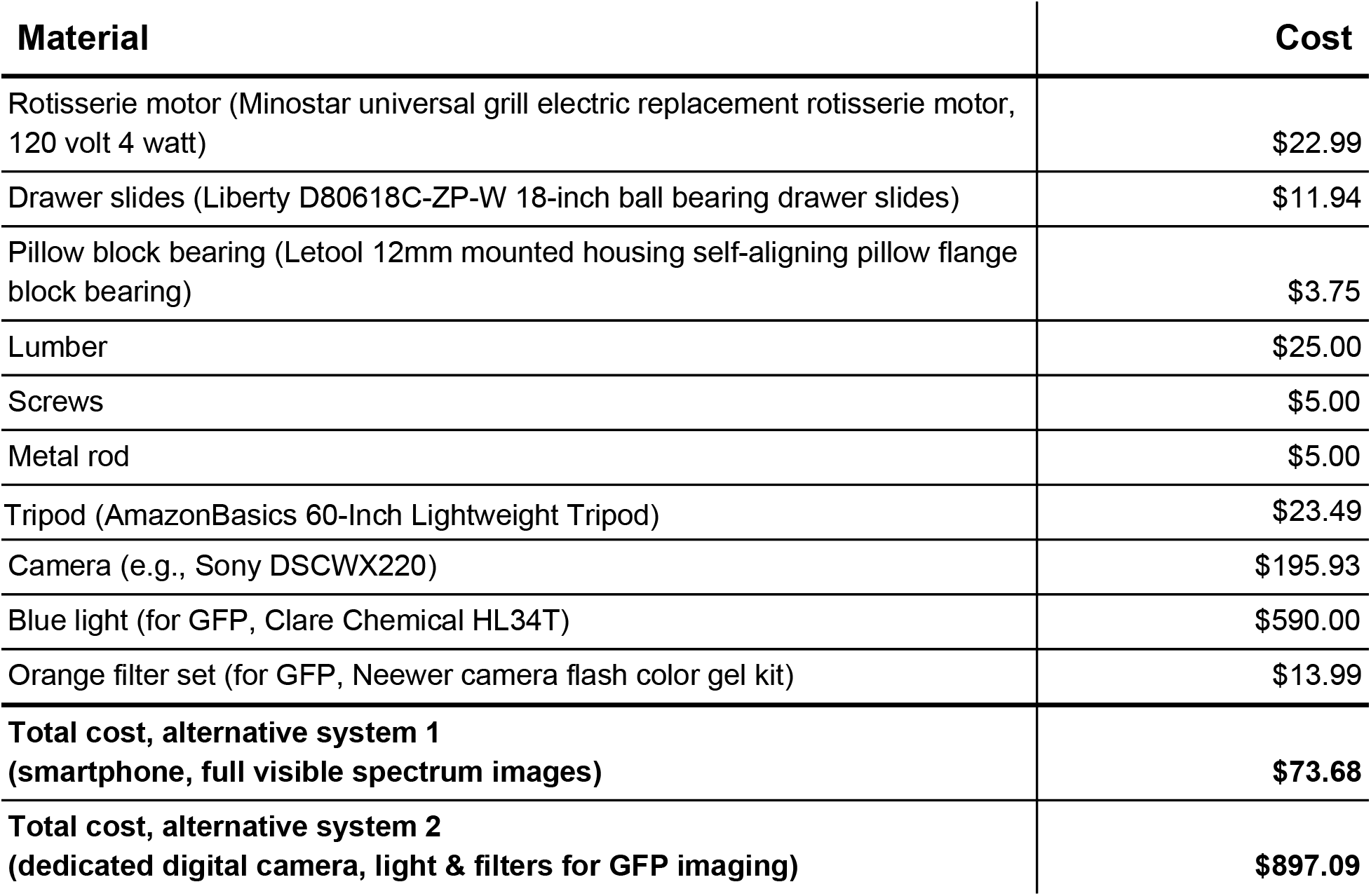
Materials and costs for scanner construction.

| **Material** | **Cost** |
| --- | --- |
| Rotisserie motor (Minostar universal grill electric replacement rotisserie motor, 120 volt 4 watt) | \$22.99 |
| Drawer slides (Liberty D80618C-ZP-W 18-inch ball bearing drawer slides) | \$11.94 |
| Pillow block bearing (Letool 12mm mounted housing self-aligning pillow flange block bearing) | \$3.75 |
| Lumber | \$25.00 |
| Screws | \$5.00 |
| Metal rod | \$5.00 |
| Tripod (AmazonBasics 60-Inch Lightweight Tripod) | \$23.49 |
| Camera (e.g., Sony DSCWX220) | \$195.93 |
| Blue light (for GFP, Clare Chemical HL34T) | \$590.00 |
| Orange filter set (for GFP, Neewer camera flash color gel kit) | \$13.99 |
| **Total cost, alternative system 1**<br>**(smartphone, full visible spectrum images)** | **\$73.68** |
| **Total cost, alternative system 2**<br>**(dedicated digital camera, light & filters for GFP imaging)** | **\$897.09** |

**Figure 1.**
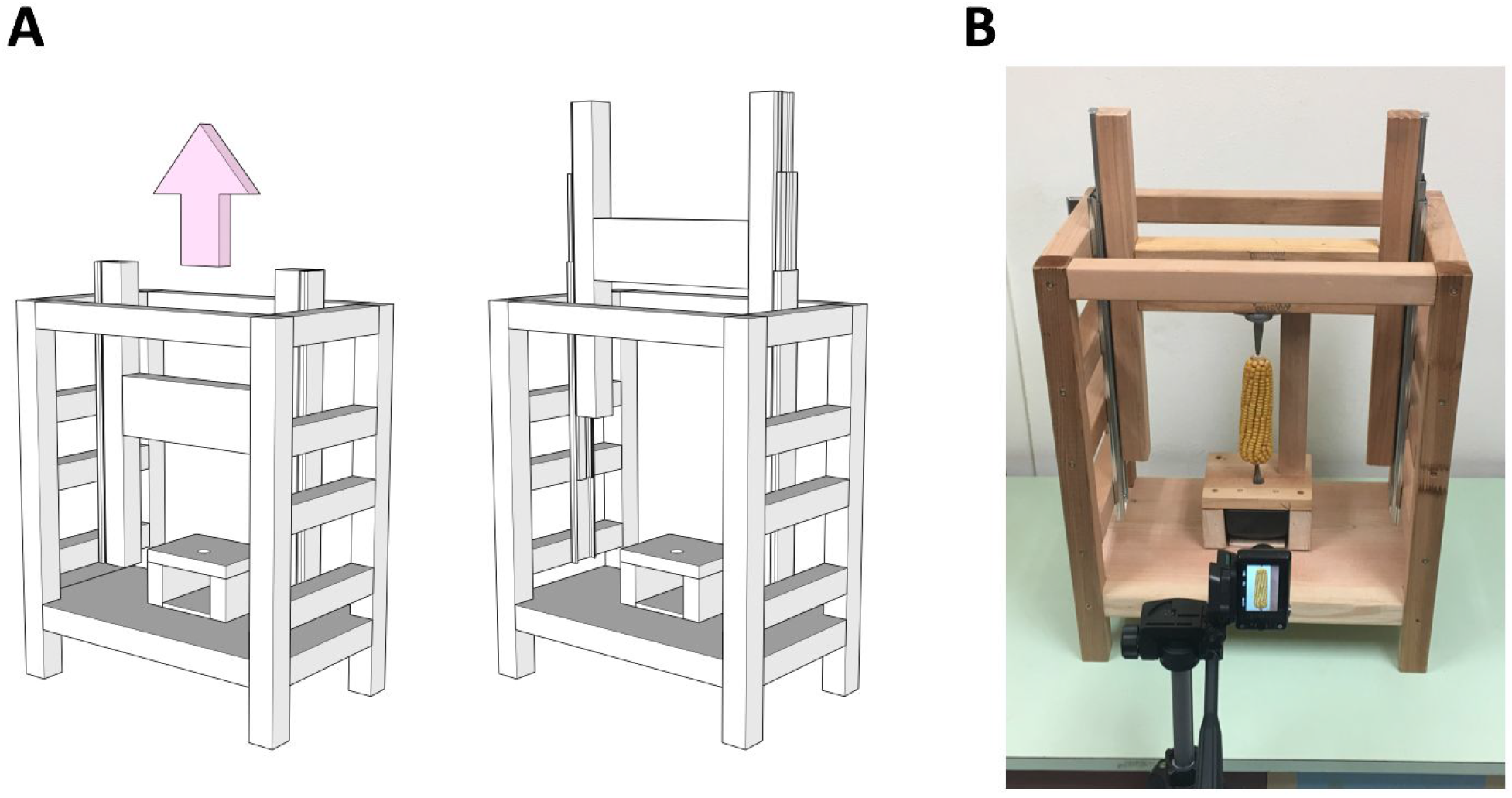
Efficient, cost-effective maize ear phenotyping with rotational scanner. **(A)** Schematics of rotational ear scanner in closed position (left) and open position (right). Full construction diagrams are available in Supplemental File 1. **(B)** Image of scanner with ear in place. Camera is positioned in front of the ear as shown, with the ear centered in the frame. A video is captured as the ear spins through one full rotation, which is then processed to project the surface of the ear onto a single flat image.

### Processing videos into flat ear projections

The output of the scanner is a video of the rotating ear. This video could be directly quantified, but we found a ‘flat’ image projection most useful for visualizing the entire surface of the ear, as well as for quantifying the distribution of seed markers. To produce this projection, videos were first uploaded to a local computer and annotated with identifying metadata (Figure 2A). Videos were then transferred to a high-performance computing cluster to be processed for generation of the projections; while this video processing step is more efficient on a computing cluster, it can alternatively be completed on a local computer. After processing, the resulting flat images were transferred back to a local computer for assessment and quantification. Video processing consisted of three steps (Figure 2B). In the first, frames were extracted from the video into separate images using the command line utility FFmpeg. Next, images were cropped to the center horizontal row of pixels using the command line utility ImageMagick. Finally, all rows of pixels, one from each frame, were appended sequentially, resulting in the final image.

**Figure 2.**
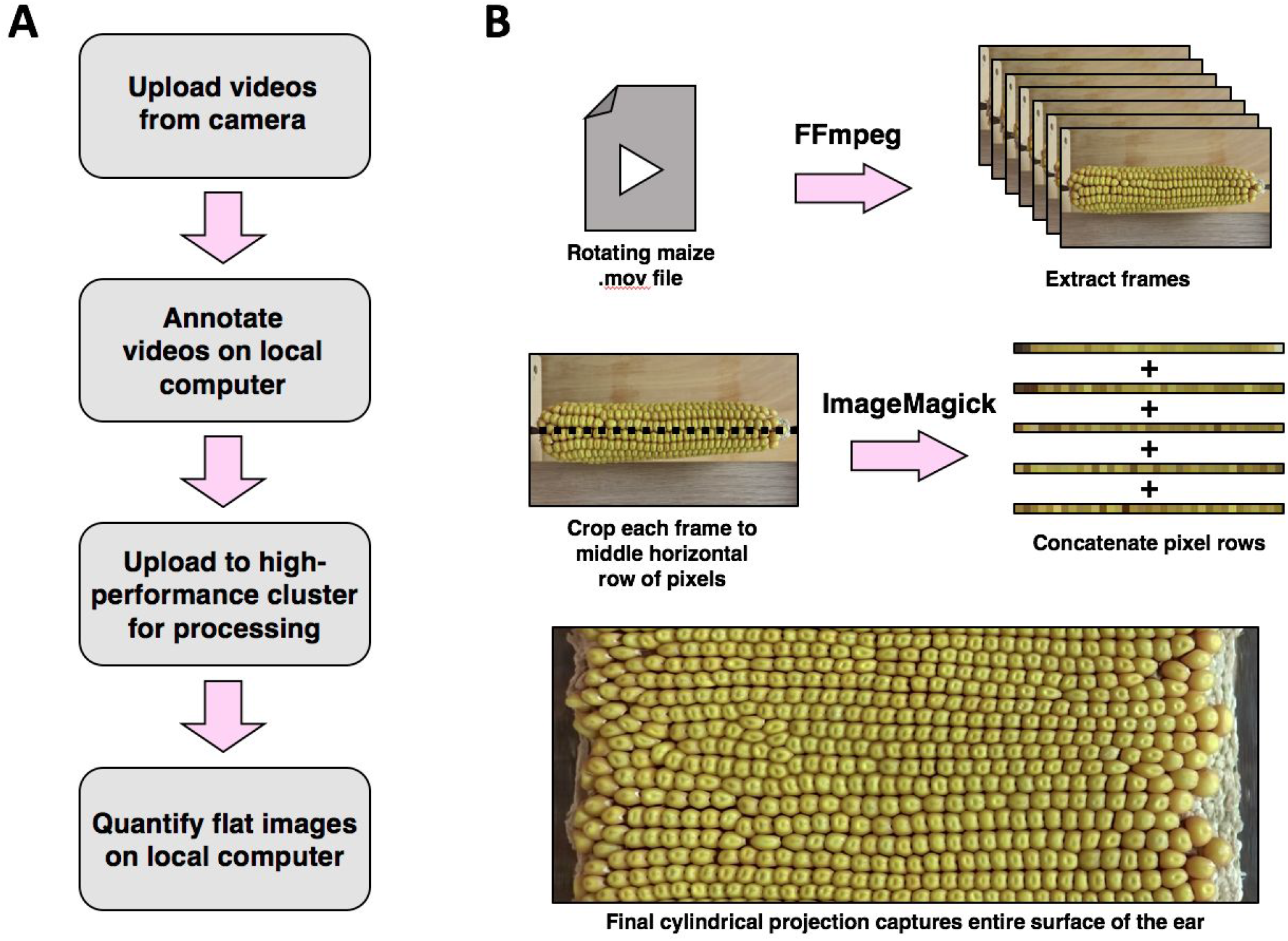
Processing videos into flat ear projections. **(A)** Image annotation and processing workflow. Videos are annotated on a local computer, followed by generation of the projection (‘flat image’) via processing either on the same computer or on a high-performance computing cluster for speed improvements. Flat images are small in size, and can be quantified on a local computer. **(B)** The video flattening process begins by extracting individual frames using FFmpeg. After frames are extracted, each frame is cropped to the middle horizontal row of pixels using the command line utility ImageMagick. The resulting collection of pixel rows, one per frame, is then concatenated into a single image depicting the entire surface of the ear.

### Example images, manual quantification, and test case

Our scanner was tested using a variety of maize ears representing several widely used seed markers (Figure 3A). Both anthocyanin *(c1)* and fluorescent *(Ds-GFP)* seed markers were easily distinguishable in the final images, as well as other markers such as *bt1, a2,* and *pr1.* Color and fluorescent seed markers were manually quantified on the digital projections using the FIJI distribution of ImageJ (Figure 3B). Using this approach, annotation of an entire ear could be completed in 5 to 10 minutes, depending on the size of the ear and the relative experience level of the annotator. In addition to producing total quantities of each seed marker, this process results in coordinates for each annotated seed, which can be further analyzed if desired. Manual annotations of scanner images in ImageJ were compared to manually counting the seeds on the ear (Figure 3C). We observed a strong correlation between these two methods (R^2^ > 0.999), validating our scanner method. To test the utility of the maize ear scanner, we scanned and quantified over 400 ears with marker-linked mutations in >50 genes. With these methods, we were able to detect weak but significant transmission defects (~45% transmission of a marker-linked mutation) for a number of mutant alleles, using both anthocyanin and GFP seed markers.

**Figure 3.**
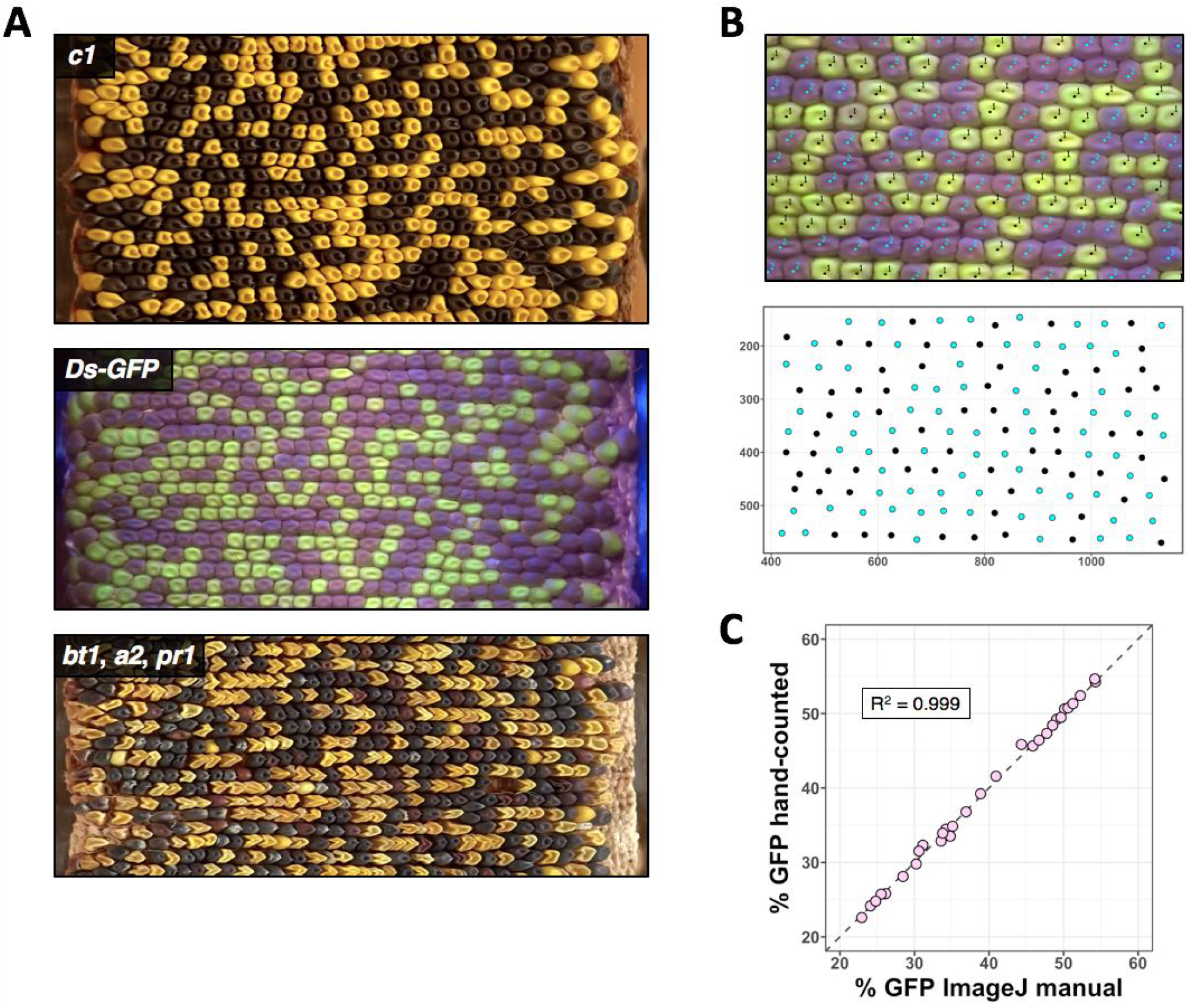
Examples of ear surface projections and quantification using ImageJ. **(A)** Representative ear projections demonstrating ease of visibility and tracking for several widely-used maize seed markers. From top to bottom: *c1; Ds-GFP; bt1/a2/pr1* (linked markers on chromosome 5). **(B)** Seed phenotypes were manually annotated on the ear image using the ImageJ Cell Counter plugin (detail top), allowing quantification of relative transmission of each marker. In this process, seed location as well as marker identities are recorded in an output xml file, allowing for optional downstream analysis of seed distributions (bottom). **(C)** Comparison of manually counting seeds on ears vs manually counting seeds from images.

## Discussion

Maize ears encode a vast amount of information. In agriculture, they provide insights into the value of a crop through yield and seed quality. In basic research, they open a window to molecular biology through mutant phenotypes and the transmission of seed markers. For example, assessing transmission rates of marked mutations on the maize ear (with seed populations of up to 600 progeny from a single cross) can generate statistically powerful data, which can then provide biological evidence for gene function during the haploid gametophyte phase. Our goal was to develop a methodology to capture some of this information via digital imaging to facilitate downstream quantitative analyses. Through capturing a video of a rotating ear, a flat image of the surface of the ear can be created that enables standardized, replicable phenotyping of seed marker distribution, as well as providing a permanent digital record of ears. Our scanner is fast, cost-effective, and capable of bringing digital image phenotyping to any lab interested in maize ears, dramatically scaling up the amount of quantitative data that can be feasibly generated.

While our scanner provides very useful phenotyping data, it has some notable limitations. Cylindrical projections are a convenient way of visualizing the entire surface of an ear in a single image. However, because maize ears are not perfect cylinders, the projections distort regions of the ear that are not cylindrical, typically the top and bottom, resulting in seeds that appear larger than those in the middle of the ear (Figure 3A). Because of these distortions, measuring qualities like seed and ear dimensions can be challenging. While approximate values for these metrics can be calculated, in the future more precise measurements could potentially use the source video as input to model the ear in three dimensions. In addition, curved ears become highly distorted when scanned using this method. Thus, we have limited the use of the projections to relatively straight, uniform-thickness ears.

With low cost as a primary goal, our scanner design has room for optimizations and improvements. Among these is better integration of the mechanisms for ear rotation, video capture, and processing. One improvement would be to drive both a configurable motor and a camera from a simple computer, such as a Raspberry Pi. Although these alterations would add cost and complexity to the process, the efficiency gains would likely offset these sacrifices.

Capturing video data from ears produces a lasting record of experiments. These data can be used in a variety of ways, such as measuring patterns of seed distribution, quantifying empty space on the ear, and recording other phenotypes such as abnormal or aborted seeds. Ultimately, recording ear data future-proofs experiments, which may benefit from yet-undeveloped methods of quantification. One future quantification method is automated seed counting. Hand annotation of seeds on ear projections using ImageJ is significantly faster than marking seeds on the ear, but remains a time-consuming and tedious process. However, the resulting data can be used to train machine-learning models to identify seeds. As these models are developed, they are likely to dramatically accelerate the phenotyping process.

## Materials and Methods

### Building the maize ear scanner

The maize ear scanner was built from dimensional lumber and widely available parts. For detailed plans and three-dimensional models, see Supplemental File 1. The base of the scanner was built from a nominal 2×12 (38×286 mm) fir board, while the frame of the scanner was built from nominal 2×2 (38×38 mm) cedar boards. Boards were fastened together with screws. Strict adherence to materials and exact dimensions of the scanner frame is not necessary, as long as the scanner is structurally sound and large enough to accommodate ears of varying sizes.

To rotate the maize ear, a standard rotisserie motor (Minostar universal grill electric replacement rotisserie motor, 120 volt 4 watt) was attached to the base of the scanner by way of a wood enclosure. Rotisserie motors are widely available and require no specialized knowledge of electronics to use. If desired, the efficiency of the scanner could be improved by using a customized motor. The rotation speed of the rotisserie motor used in this scanner (~2 RPM) was slightly slower than optimal. A faster rotation speed could improve the overall scanning time, which is ultimately dependent on the frame rate of the camera. The lower portion of the ear was fastened to the rotisserie motor using a 5/16” (8 mm) steel rod, which can be easily removed from the motor when switching ears. The top of the steel rod was ground to a flattened point with a bench grinder to allow it to be inserted into the pith at the center of the base of the ear.

The top of the ear was held in place with an adjustable assembly constructed from a nominal 2×4 board (38×89 mm) fastened to drawer slides (Liberty D80618C-ZP-W 18-inch ball bearing drawer slides) on either side of the scanner frame (Supplemental File 1). In the center of the 2×4, facing down towards the top of the ear, is a steel pin mounted on a pillow block bearing (Letool 12mm mounted housing self-aligning pillow flange block bearing). The steel pin (12mm) was sharpened to a point to penetrate the top of the ear as it is lowered, temporarily holding it in place as the ear is rotated during scanning. Because the pin can be moved up and down on the drawer slides, a variety of ear sizes can be accommodated in the scanner.

For full spectrum visible light images, ambient lighting was used. To capture GFP fluorescence, a blue light (Clare Chemical HL34T) was used to illuminate the ear. An orange filter (Neewer camera flash color gel kit) was placed in front of the camera lens to partially filter out non-GFP wavelengths. More cost effective blue light illumination is possible (e.g. Wayllshine blue LED flashlight, $9.00), however we found achieving sufficient brightness to be challenging without the higher power LED light.

### Workflow for scanning an ear

The scanning process begins by trimming the top and bottom of the ear to expose the central pith. The bottom pin is then inserted into the bottom of the ear, after which the pin with ear attached is placed in the rotisserie motor. The top of the ear is secured by lowering the top pin into the pith at the top of the ear. After turning on the rotisserie motor, a video is captured by the camera that encompasses at least one complete rotation of the ear. The ear can then be removed from the scanner, and the next ear scanned. A detailed, illustrated protocol for scanning ears with the maize ear scanner using ears with GFP seed markers and a Sony DSCWX220 can be found in Supplemental File 2.

### Creating flat images

After videos are imported to a computer, they are processed to flat images. Frames are first extracted from videos to png formatted images using the command line utility FFmpeg (https://ffmpeg.org) with default options (ffmpeg -i ./”$file” -threads 4 ./maize_processing_folder/output_%04d.png). These images are then cropped to the central row of pixels using the command line utility ImageMagick (https://imagemagick.org/) (mogrify -verbose -crop 1920×1+0+540 +repage ./maize_processing_folder/*.png). The collection of single pixel row images is then appended in sequential order (convert -verbose -append +repage ./maize_processing_folder/*.png ./”$name.png”). Finally, the image is rotated and cropped (mogrify -rotate “180” +repage ./”$name.png”; mogrify -crop 1920×746+0+40 +repage ./”$name.png”). We chose the convention of a horizontal flattened image with the top of the ear to the right and the bottom of the ear to the left. Because our videos were captured vertically, a rotation was required after appending the individual frames. Depending on the users video capture orientation and desired final image orientation, some modification may be necessary. The dimensions of the final crop of the image depend both on the rotation speed and the video frame rate. It is not necessary to capture one exact rotation, as long as the video captures at least one rotation. If the ears rotate at a consistent speed, one full rotation will always be the same number of frames. Because each frame becomes one pixel in height, the image can be cropped to a height corresponding to the number of frames that encompass one rotation. Because rotation speeds and frame rates vary, it is recommended that the user records the dimensions of an ear and compares those dimensions to the final output to ensure that there is no distortion. The final crop can be adjusted to address any possible distortion. For very high frame rates, the FFmpeg frame extraction rate can also be adjusted using the -vf option and adjusting the playback frames per second (e.g. to extract 10 frames per second: -vf fps=10). A GitHub repository containing the script used to create flat images from videos is located at https://github.com/fowler-lab-osu/flatten_all_videos_in_pwd.

### Quantifying seeds using flat images

Seeds were quantified from flat ear images using the Cell Counter plugin of the FIJI distribution of ImageJ (Schindelin et al., 2012). Ears were assigned counter types to correspond to different seed markers, after which seeds on ear images were located and annotated by hand. The Cell Counter plugin exports results in an xml file, which contains the coordinates and marker type of every annotated seed. This file can be processed to create a map of seed locations on the ear. A detailed protocol describing the quantification process can be found in Supplemental File 3.

## Acknowledgements

We thank O. Childress, H. Fowler, B. Galardi, B. Hamilton, R. Hartman, C. Lambert for their seed counting assistance. In addition, we thank O. Childress for protocol feedback and J. Preece for useful discussions. This work was supported by NSF grants IOS-1340050 and IOS-1832186, as well as funds from the OSU Department of Botany and Plant Pathology. We also thank the OSU Center for Genome Research and Biocomputing for providing computational infrastructure enabling this project. This invention is patent pending with Oregon State University.

## Supplemental Data

**Supplemental File 1.** Rotational ear scanner construction plans.

**Supplemental File 2.** Scanning fluorescent ears with the rotational ears scanner protocol.

**Supplemental File 3.** Quantifying seeds in flat images using ImageJ protocol.

