## Supplementary figures and images for "Custom built scanner and simple image processing pipeline enables low-cost, high-throughput phenotyping of maize ears"

### image_of_scanner.png

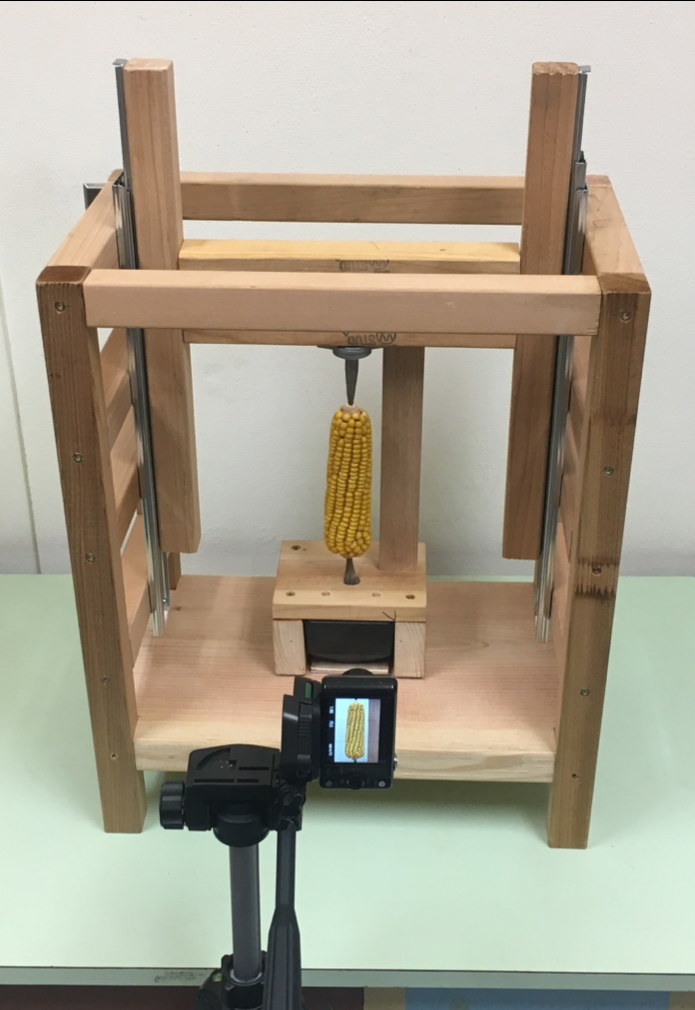

### scanner_plans_front_view.pdf

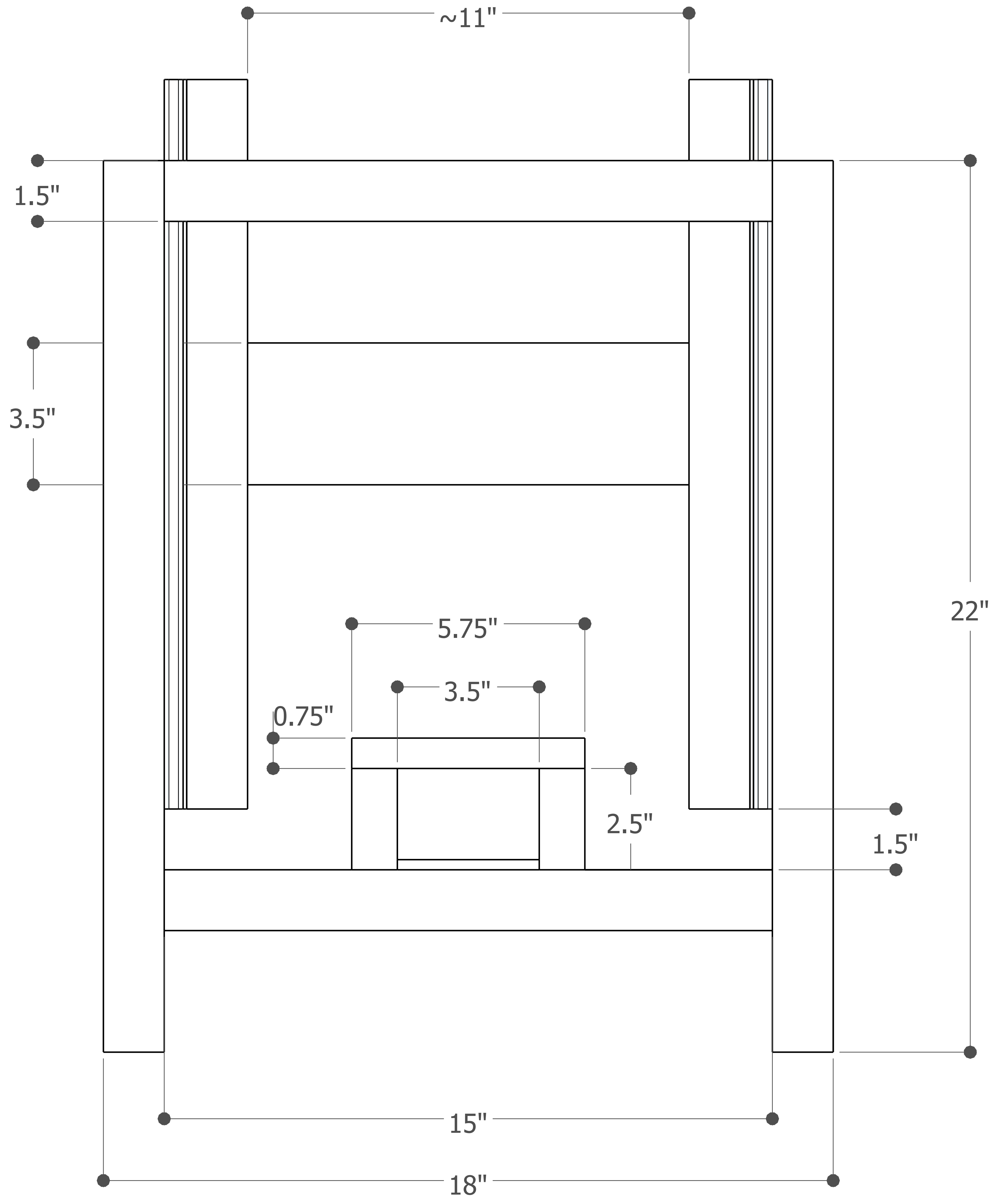

### scanner_plans_no_dimensions_closed.pdf

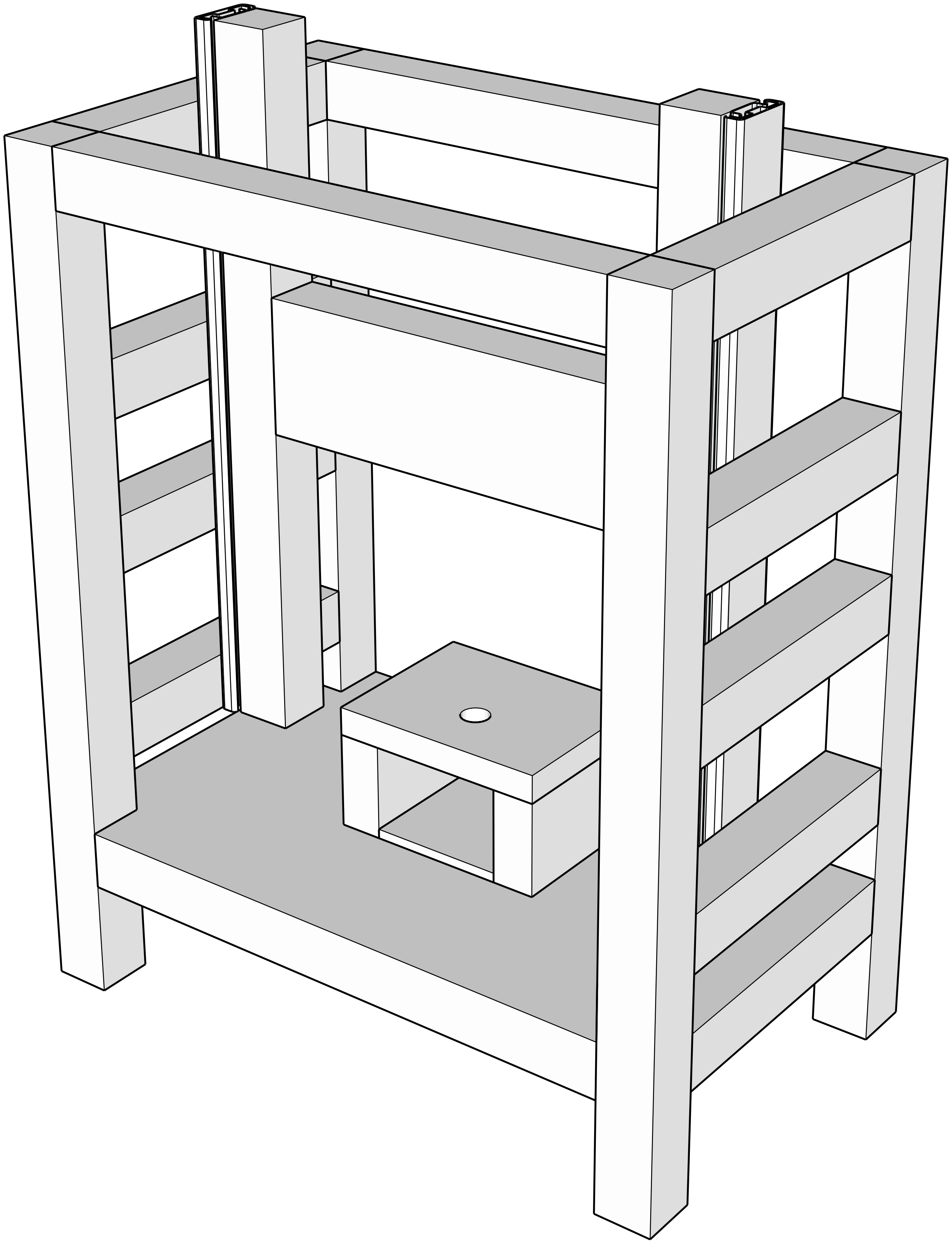

### scanner_plans_no_dimensions_open.pdf

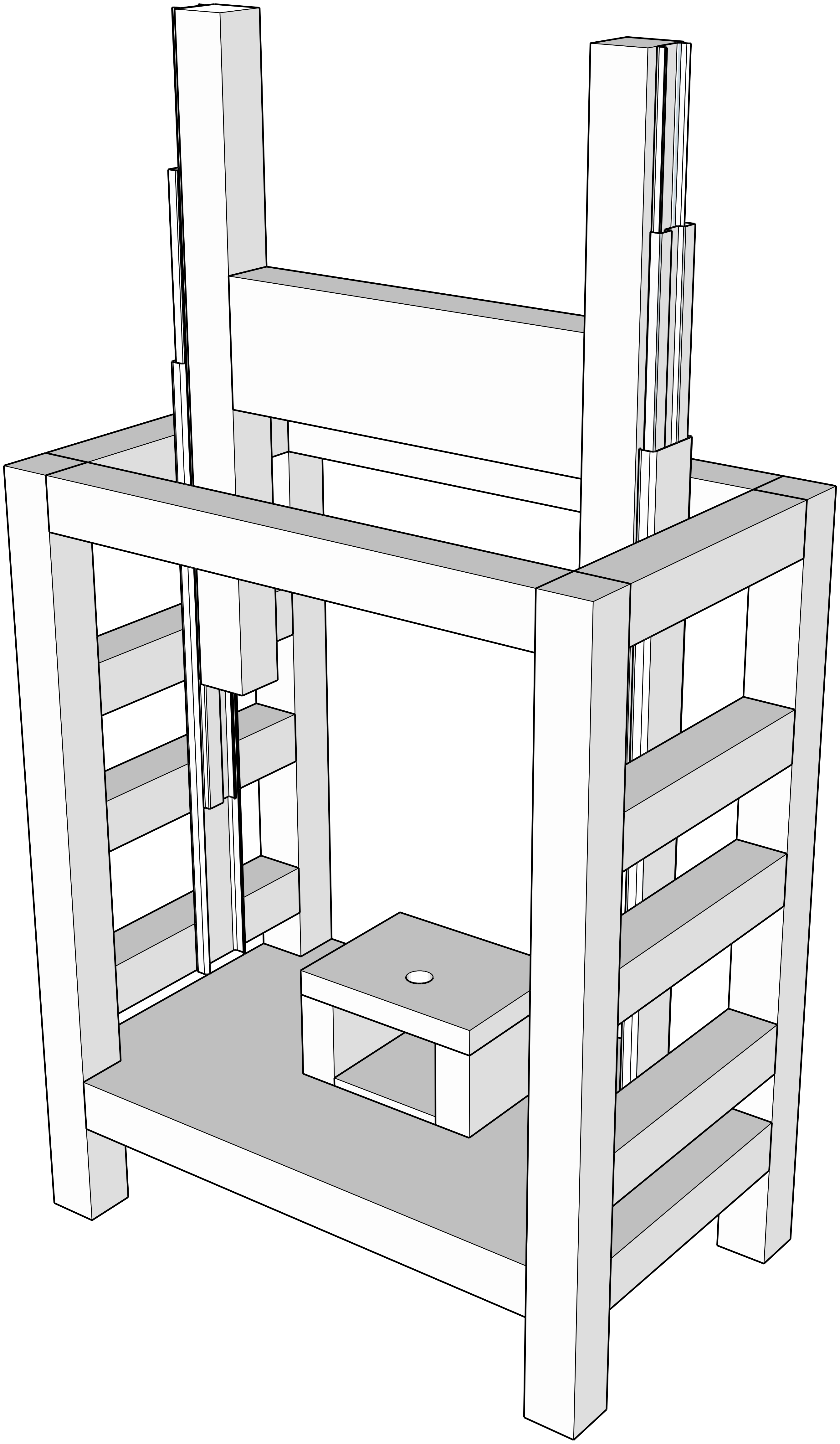

### scanner_plans_side_view.pdf

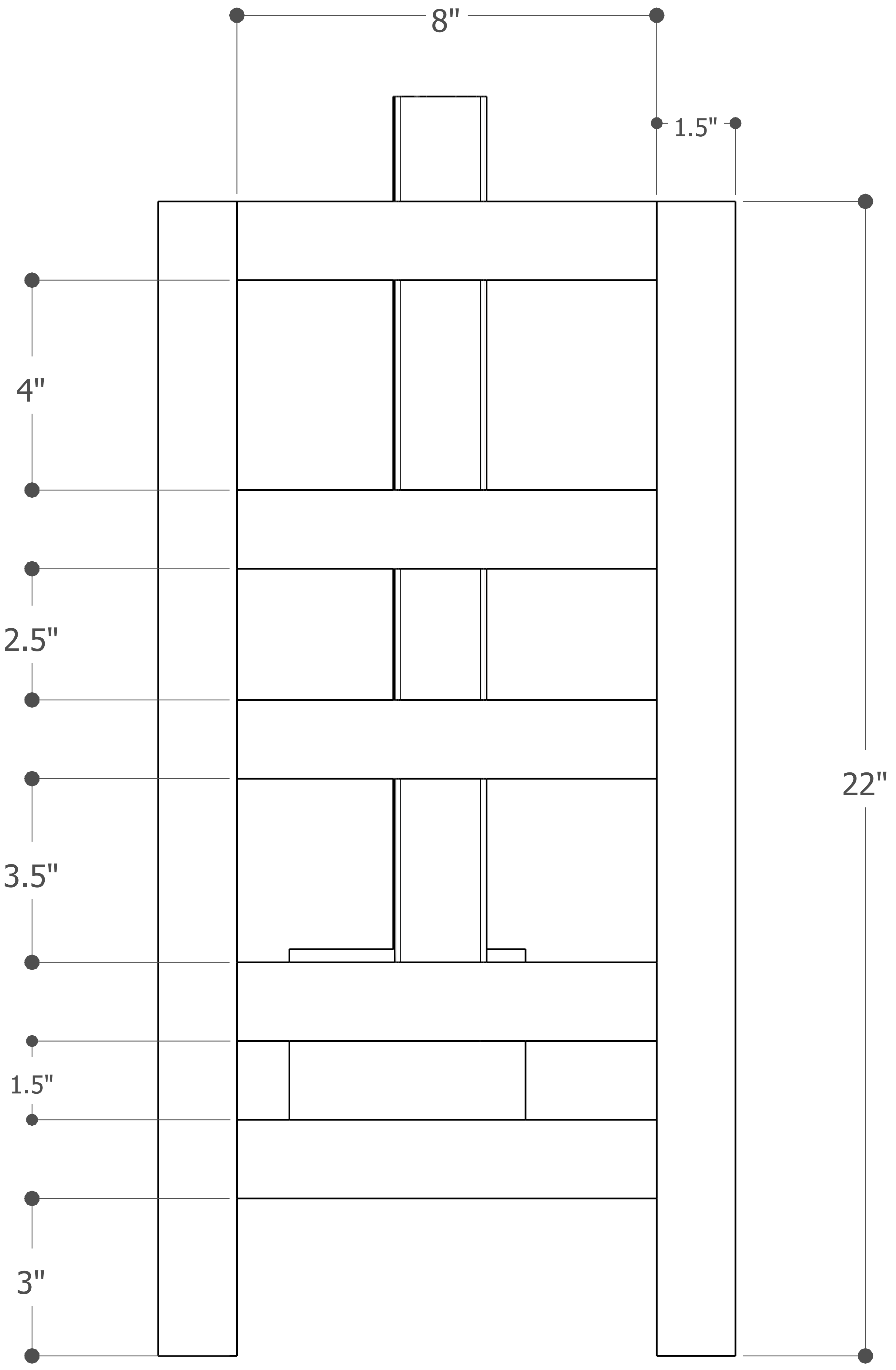

### scanner_plans_top_view.pdf

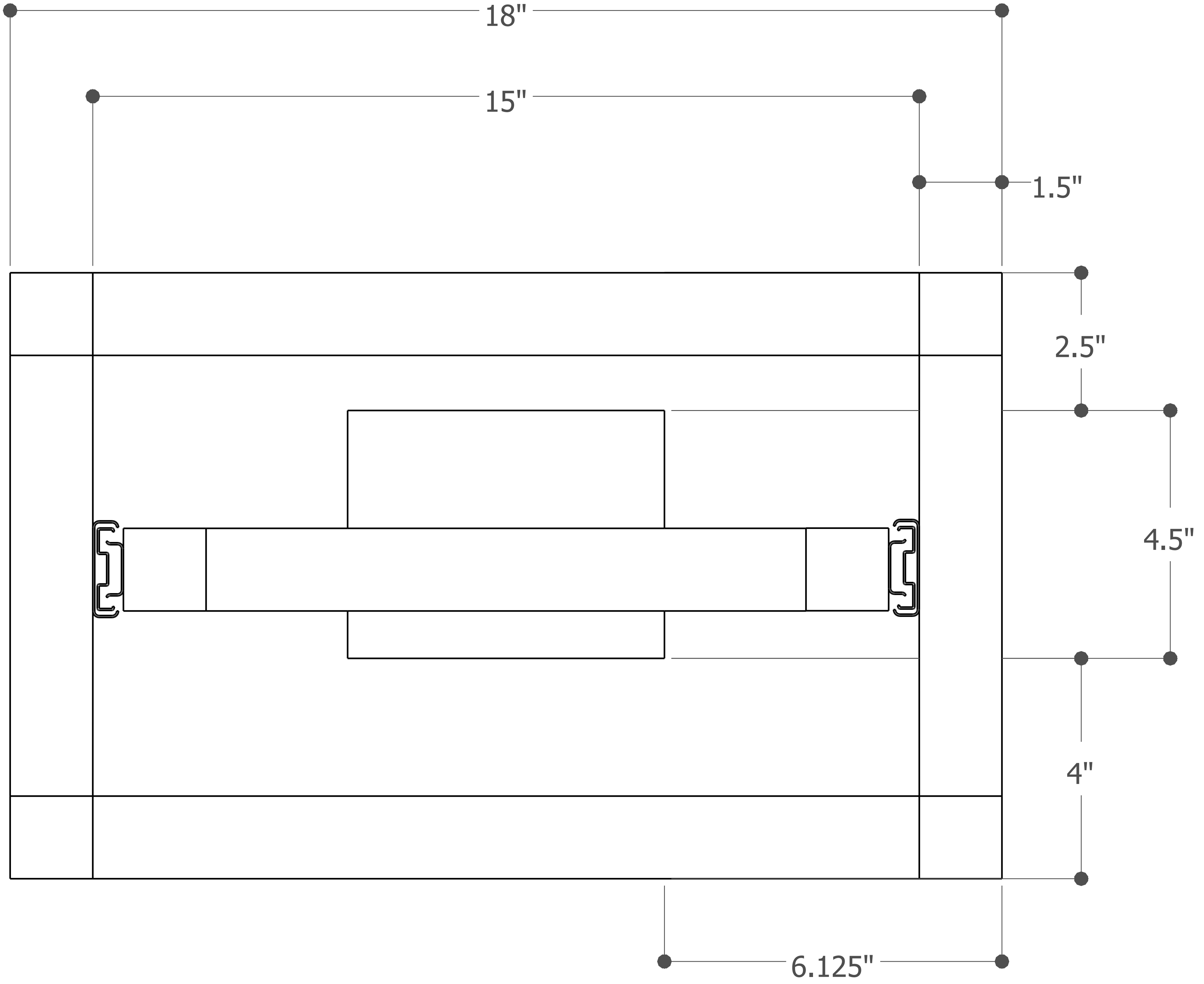

### scanner_plans_with_dimensions.pdf

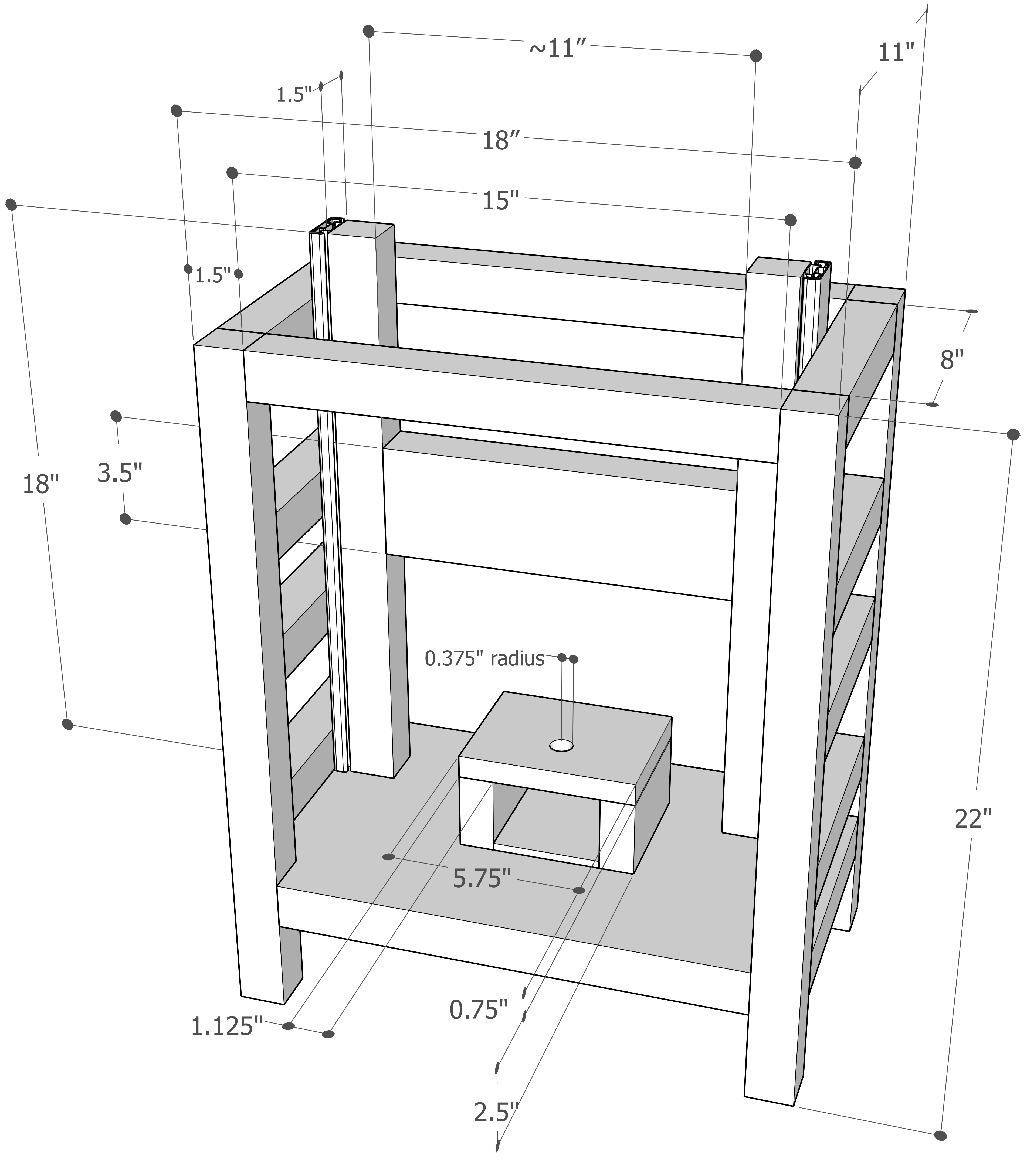
