## Supplemental File 2 for "Custom built scanner and simple image processing pipeline enables low-cost, high-throughput phenotyping of maize ears"

### Scanning fluorescent ears with the maize ear scanner

*This protocol describes the process of scanning maize ears with green fluorescent kernels (dsGFP). A video is created of the rotating ear, which is then flattened into an image containing the entire surface of the ear.*

1. Cut excess material off the top and bottom of the ear. The pith in the center of the ear should be exposed on both the top and bottom of the ear. Remove silks.

**Before:**

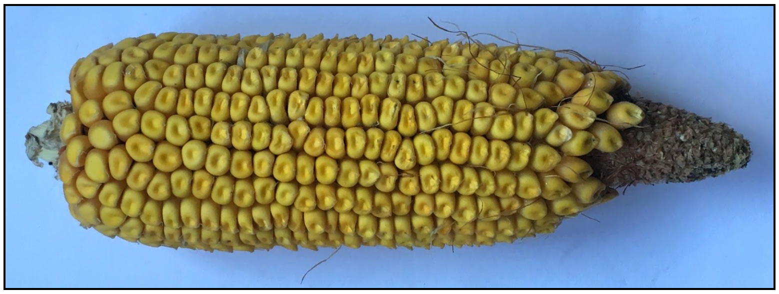

**After:**

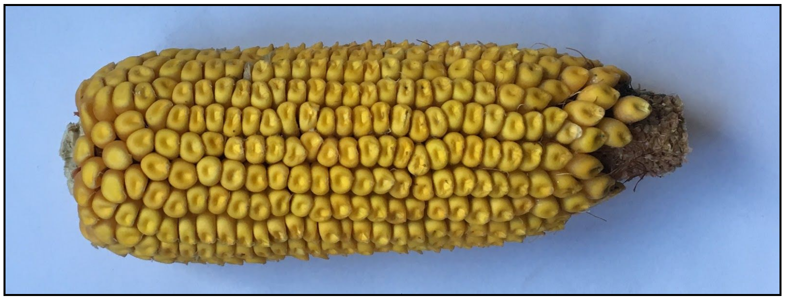

2. Insert bottom skewer into the pith on the base of the ear as shown, attempt to insert it as centered as possible. If you need to push hard to get it into the ear, use gloves.

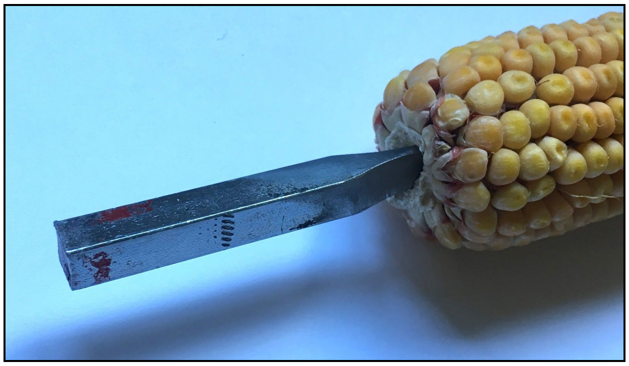

6. Raise top skewer and place bottom skewer in rotisserie.
7. Lower top skewer onto center of top of ear. Use caution when kernels go all the way to the top of the ear; it's possible to push too hard on the top skewer and forcibly dislodge several seeds. If kernels go all the way to the top of an ear, the top 10-20 kernels can be removed to make room for the top skewer. After removing the top kernels, trim the top of the ear to make a good position for the skewer.

**Ear in place and ready to scan:**

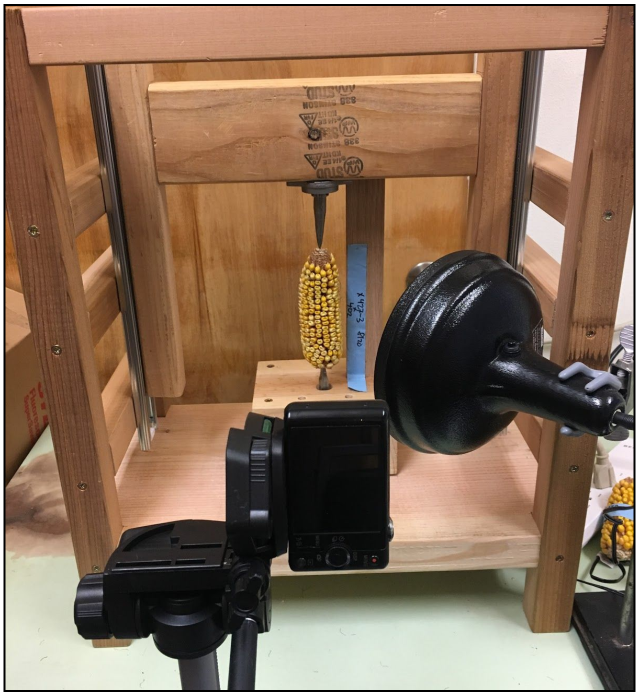

8. Turn off the lights in the scanning room.
9. Adjust the blue light so that it shines perpendicular to the axis of ear rotation from the right side. The light should be vertically centered at the halfway point of the ear. In

addition, place the light as close to the camera as possible, without the light obscuring any of the kernels.

**Optimal light placement:**

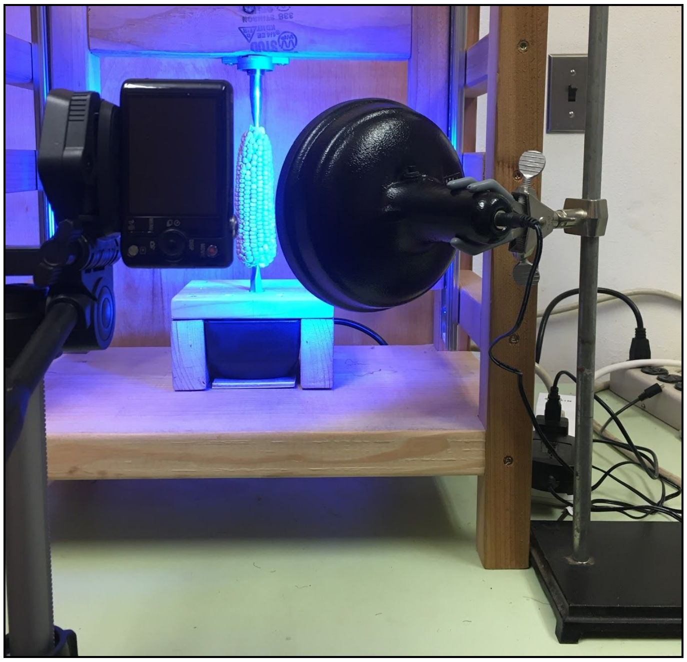

10. Turn on the camera. For the camera we used (Sony DSCWX220), adjust settings as follows: "High Sensitivity Movie" mode (Menu -> Camera Settings -> Movie -> High Sensitivity), 60i 17M(FH) resolution (Menu -> Camera Settings -> Record Setting -> 60i 17M(FH)), with image stabilization turned off (Menu -> Camera Settings -> SteadyShot -> Standard).
11. Ensure that the orange filter has been placed in front of the camera lense, and that glare from the light is minimized.
12. Align camera vertically as shown in the image above.
13. Adjust camera height on tripod. The camera lense should be centered on the halfway point of the ear.
14. Zoom in until the ear nearly fills the frame.
15. Place the ear in the exact center of the frame using the reference lines in the camera viewfinder. It is essential that the long dimension of the ear is exactly centered.

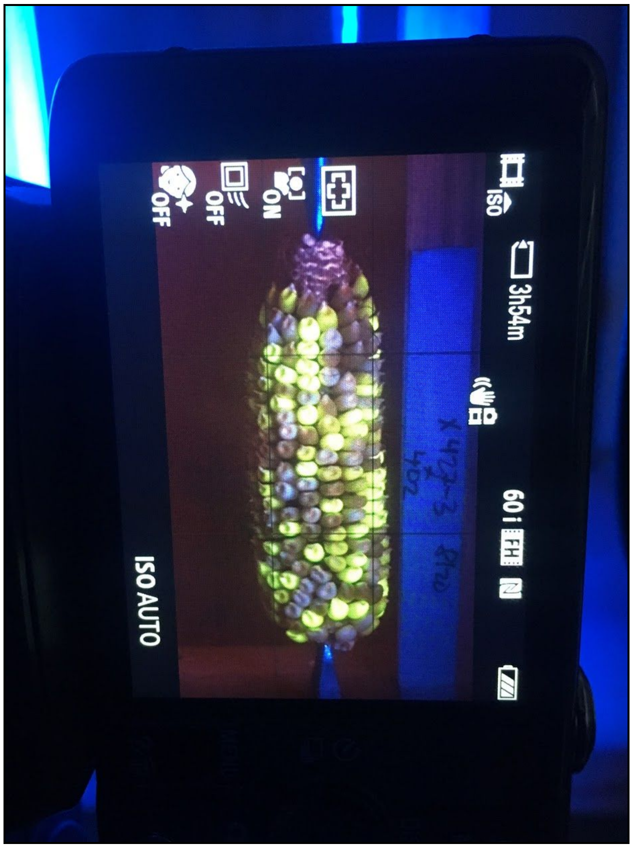


16. Turn on the rotisserie motor. Capture a 30 second long video of the rotating ear. Longer than 30 seconds is ok, but less than 30 seconds is not (this will vary depending on your rotisserie motor's speed). Ensure that the camera remains completely steady during the video capture; if the tripod shakes, retake the video. Make sure that the ear is rotating before the video begins and after it is finished. Check that the fluorescent kernels aren't overexposed (there's no way that I know of to manually adjust the exposure in a video on this camera, but rotating the blue light so that it hits the ear on an angle works well).

**Overexposed:**

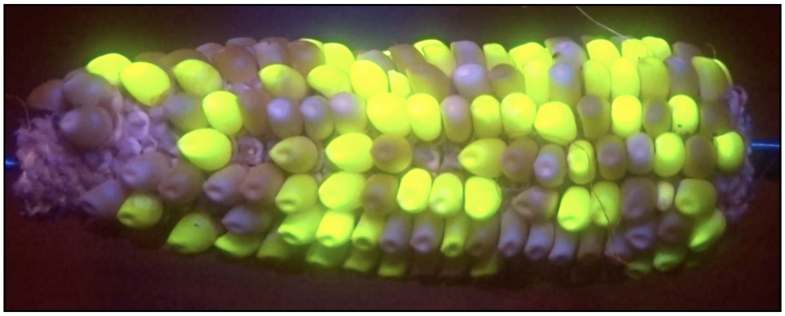

**Correct exposure:**

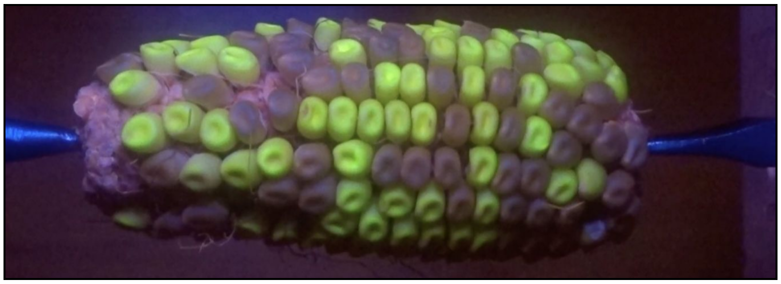

17. Remove ear from scanner, repeat for all ears.
