## Supplemental File 3 for "Custom built scanner and simple image processing pipeline enables low-cost, high-throughput phenotyping of maize ears"

### Quantifying seeds in flat images using ImageJ

*This protocol describes the process of using ImageJ to count seeds from flat images of fluorescent ears (dsGFP). An ImageJ distribution named FIJI was used for this protocol, version 2.0.0-rc-68/1.52h on MacOS 10.12.6.*

1. If it's not already installed, download the FIJI distribution of ImageJ.
2. Open your image in ImageJ.
3. We will be using the "Cell Counter" plugin to analyze our image. To open Cell Counter, navigate to Plugins -> Analyze -> Cell Counter -> Cell Counter.

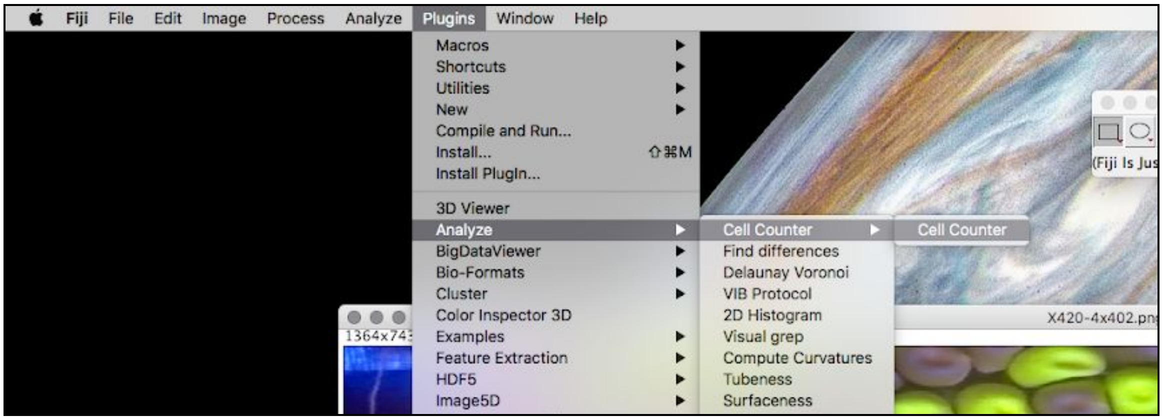

4. Selecting Cell Counter will open a new window. Press the "Initialize" button in the new window. This creates a "Counter Window," which basically reopens your image. To increase the size of the image, select Image -> Zoom -> Maximize.

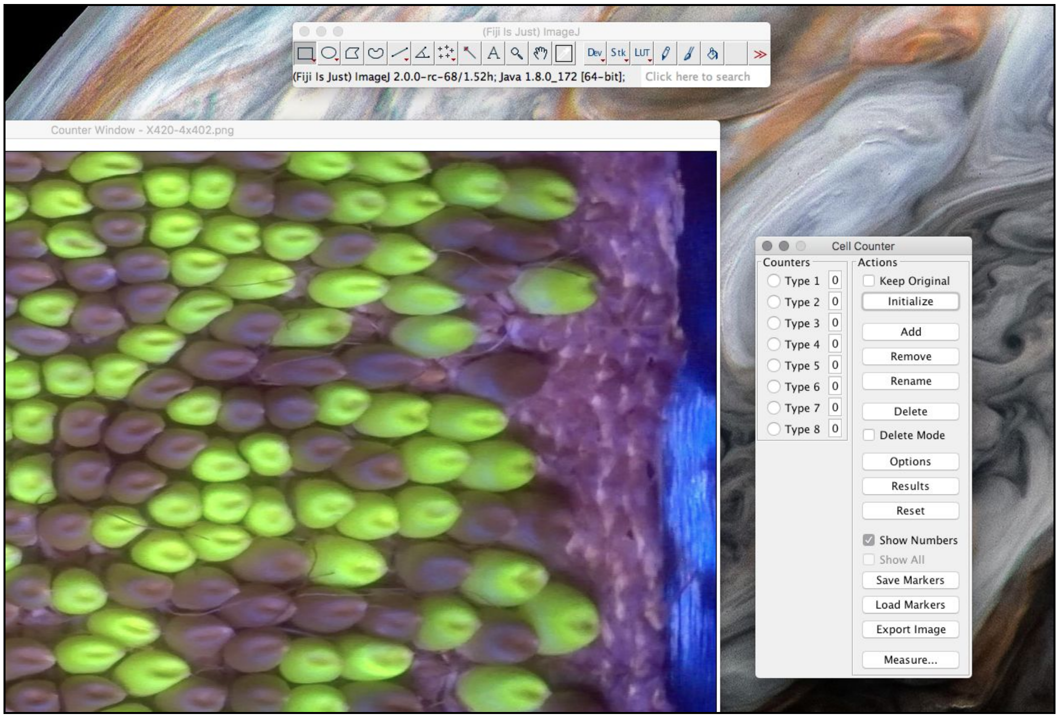


5. For this protocol, we will count the fluorescent seeds using the "Type 1" counters, and the non-fluorescent seeds using the "Type 2" counters. To begin counting fluorescent seeds, click the button to the left of Type 1.

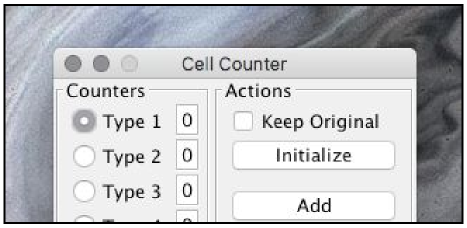


6. In the counter window, click the center of each fluorescent seed. A "1" should appear in each clicked position. If individual seeds are split between the bottom and top of the image, count the larger portion of the seed to avoid double counting. Note that the running tally of total marked seeds can be found in the box to the right of the Type 1 counter selection in the Cell Counter window.

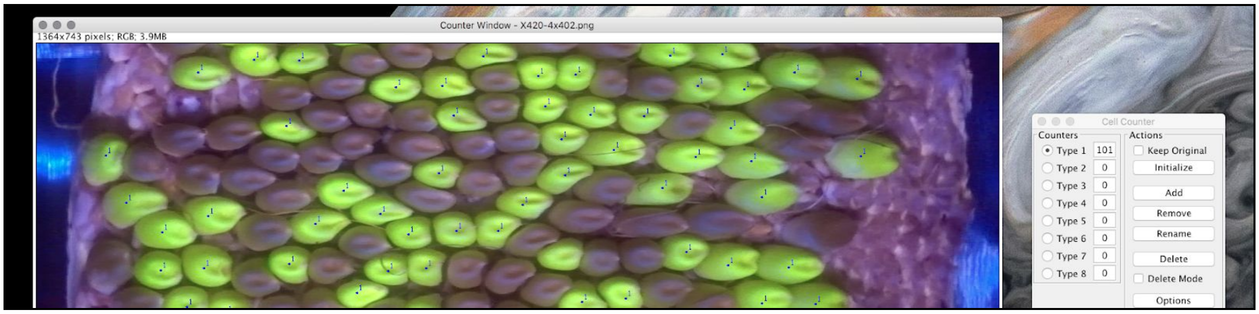


7. If you mistakenly click on the image where you don't intend to, click the delete mode box in the cell counter window. Click near the accidental point to remove it. Note: to delete Type 1 counters, Type 1 must be selected in the Cell Counter window. If another type is selected, it will delete the nearest marker for the other type.
8. Once all fluorescent seeds are marked, proceed to counting the non-fluorescent seeds. These seeds will be marked with Type 2 counters, so click the circle to the left of Type 2 in the cell counter window.

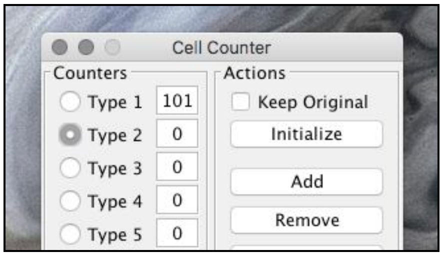


9. Count all non-fluorescent seeds in the same way as fluorescent seeds. A small "2" should appear on each non-fluorescent seed you click.

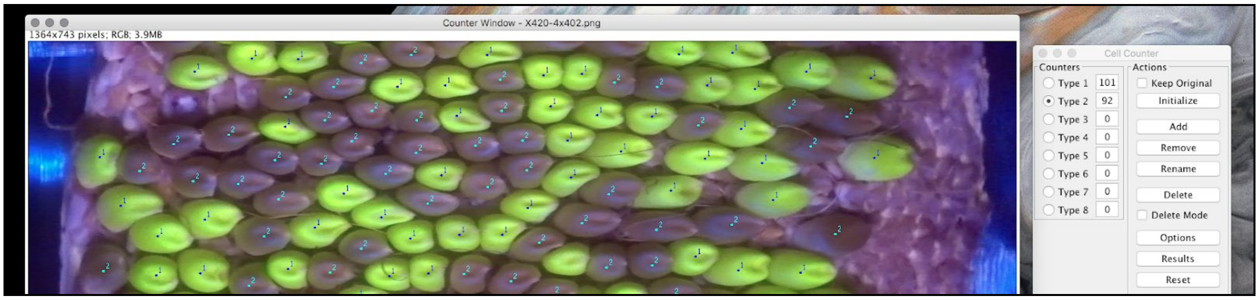


10. Unclear seeds can be marked with Type 3 counters in the same way as previously described.
11. Total numbers of each seed type can be recorded after annotations are complete.
12. To save counter positions, click the "Save Markers" button in the Cell Counter window. Counters can be saved at any point in the process, overwrite the previous file if you save

more than once. Counter positions will be saved in .xml format, which can be parsed for downstream analysis.

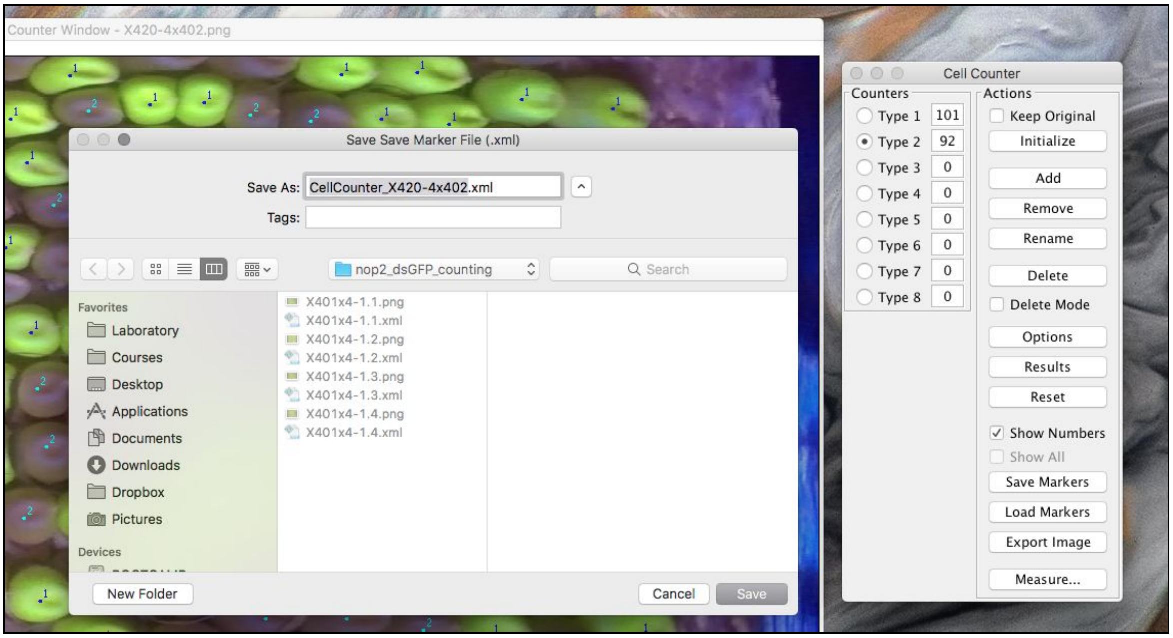


13. Before counting the next image, close both the current image and the cell counter window.
